# Topological analysis of the type 3 secretion system translocon pore protein IpaC following its native delivery to the plasma membrane during infection

**DOI:** 10.1101/605865

**Authors:** Brian C. Russo, Jeffrey K. Duncan, Marcia B. Goldberg

**Author notes:** To whom correspondence should be addressed: Marcia B. Goldberg, Division of Infectious Diseases, Massachusetts General Hospital, 4 Blackfan Circle, Boston, MA 02115.

## Abstract

Many Gram-negative bacterial pathogens require a type 3 secretion system (T3SS) to deliver effector proteins into eukaryotic cells. Contact of the tip complex of the T3SS with a target eukaryotic cell initiates the secretion of the two bacterial proteins that assemble into the translocon pore in the plasma membrane. The translocon pore functions to regulate effector protein secretion and is the conduit for effector protein translocation across the plasma membrane. To generate insights into how the translocon pore regulates effector protein secretion, we defined the topology of the *Shigella* translocon pore protein IpaC in the plasma membrane following its native delivery by the T3SS. Using single-cysteine substitution mutagenesis and site-directed labeling with a membrane-impermeant chemical probe, we mapped residues accessible from the extracellular surface of the cell. Our data support a model in which the N-terminus of IpaC is extracellular and the C-terminus of IpaC is intracellular. These findings resolve previously conflicting data on IpaC topology that were based on non-native delivery of IpaC to membranes. *Salmonella enterica* serovar Typhimurium also requires the T3SS for effector protein delivery into eukaryotic cells. Although the sequence of IpaC is closely related to the *Salmonella* translocon pore protein SipC, the two proteins have unique functional attributes during infection. We showed a similar overall topology for SipC and IpaC and identified subtle topological differences between their transmembrane α-helixes and C-terminal regions. Together, our data suggest that topological differences among distinct translocon pore proteins may dictate organism-specific functional differences of the T3SSs during infection.

**Importance:** The type 3 secretion system (T3SS) is a nanomachine required for virulence of many bacterial pathogens that infect humans. The system delivers bacterial virulence proteins into the cytosol of human cells, where the virulence proteins promote bacterial infection. The T3SS forms a translocon pore in the membrane of target cells. This pore is the portal through which bacterial virulence proteins are delivered by the T3SS into the eukaryotic cytosol. The pore also regulates the secretion of these virulence proteins. Our work defines the topology of translocon pore proteins in their native context during infection, resolves previously conflicting reports about the topology of the *Shigella* translocon pore protein IpaC, and provides new insights into how interactions of the pore with the T3SS likely produce signals that activate secretion of virulence proteins.

## Introduction

The type 3 secretion system is a specialized nanomachine required for the virulence of more than 30 bacterial pathogens (1). The T3SS translocates bacterial virulence proteins, known as effector proteins, from the bacterial cytosol into the cytosol of eukaryotic cells. The T3SS is composed of a base that spans the two bacterial membranes (2), a needle that is anchored in the base and extends away from the bacterial surface (2,3), and a tip complex that prevents non-specific secretion (4,5). Upon contact of the tip complex with a eukaryotic cell, the T3SS secretes two bacterial proteins that embed in the plasma membrane (6), where they assemble into a hetero-oligomeric pore, known as the translocon pore (3). The translocon pore is essential for T3SS activity; it functions as a conduit through which bacterial virulence proteins (“effectors”) traverse the plasma membrane to gain access to the eukaryotic cytosol (3), and it participates in defining the timing of the secretion of these effectors by the T3SS (7).

*Shigella flexneri* causes bacterial dysentery in humans and requires a T3SS to invade into and spread among the epithelial cells of the colon. *S. flexneri* has become a model organism with which to investigate the T3SS structure and function. The *S. flexneri* translocon pore is composed of the proteins IpaB and IpaC. Interactions of IpaC with host intermediate filaments are required for the *Shigella* T3SS to stably associate with the translocon pore in a process known as docking (7). Docking is essential for the bacterium to remain adherent to the cell and to secrete virulence proteins through the T3SS (7,8). Thus, stable interactions of the T3SS needle with the pore cue the T3SS to activate secretion. Aspects of the pore accessible from the extracellular environment are predicted to interact with the needle, but it is uncertain how this interaction occurs, as the topology of IpaC in the plasma membrane is controversial. The two previous studies that investigated IpaC topology used purified recombinant IpaC. These studies showed IpaC inserting into the membrane with its N-terminus on the extracellular surface of the plasma membrane, but came to opposing conclusions about the location of the C-terminus; interactions of IpaC with artificial liposomes concluded that IpaC contained a single transmembrane α-helix with the C-terminus present in the liposome lumen (9), whereas investigation of purified IpaC incorporating into macrophage membranes concluded that IpaC contained two transmembrane α-helixes with the C-terminus accessible on the extracellular surface of the macrophage (10).

Here we defined the topology of the *Shigella* translocon protein IpaC during bacterial infection following its native delivery into the plasma membrane by the T3SS. Using single cysteine accessibility mutagenesis, we defined a topological map of IpaC showing that the amino terminal region is extracellular, that the carboxy terminal region is in the cytosol, and that a single transmembrane α-helix is present. Furthermore, to test whether this topology is conserved among IpaC homologs in other pathogens that require T3SS for virulence, the accessibility of analogous residues for the *Salmonella* pore protein SipC was tested. We found that the overall topologies of IpaC and SipC are similar. However, we observed subtle differences between the two proteins in the accessibility of the transmembrane α-helixes and the C-terminal regions that may contribute to organism-specific functional differences of these T3SSs during infection.

## Results

### Generation of single cysteine substitution derivatives of IpaC

To generate insights into how the translocon pore regulates effector protein secretion, the accessibility of specific residues within natively-delivered plasma-membrane embedded IpaC was determined. IpaC residues were selected for analysis based on a combination of previous *in silico* analysis of the putative IpaC secondary structure (7) (Fig. 1A), and since our approach involved single cysteine substitution, on previous successful mutation of particular IpaC residues (9,11) and an attempt to minimize the negative impact of the cysteine substitution on protein function by choosing alanine or serine residues for replacement.

**Figure 1:**
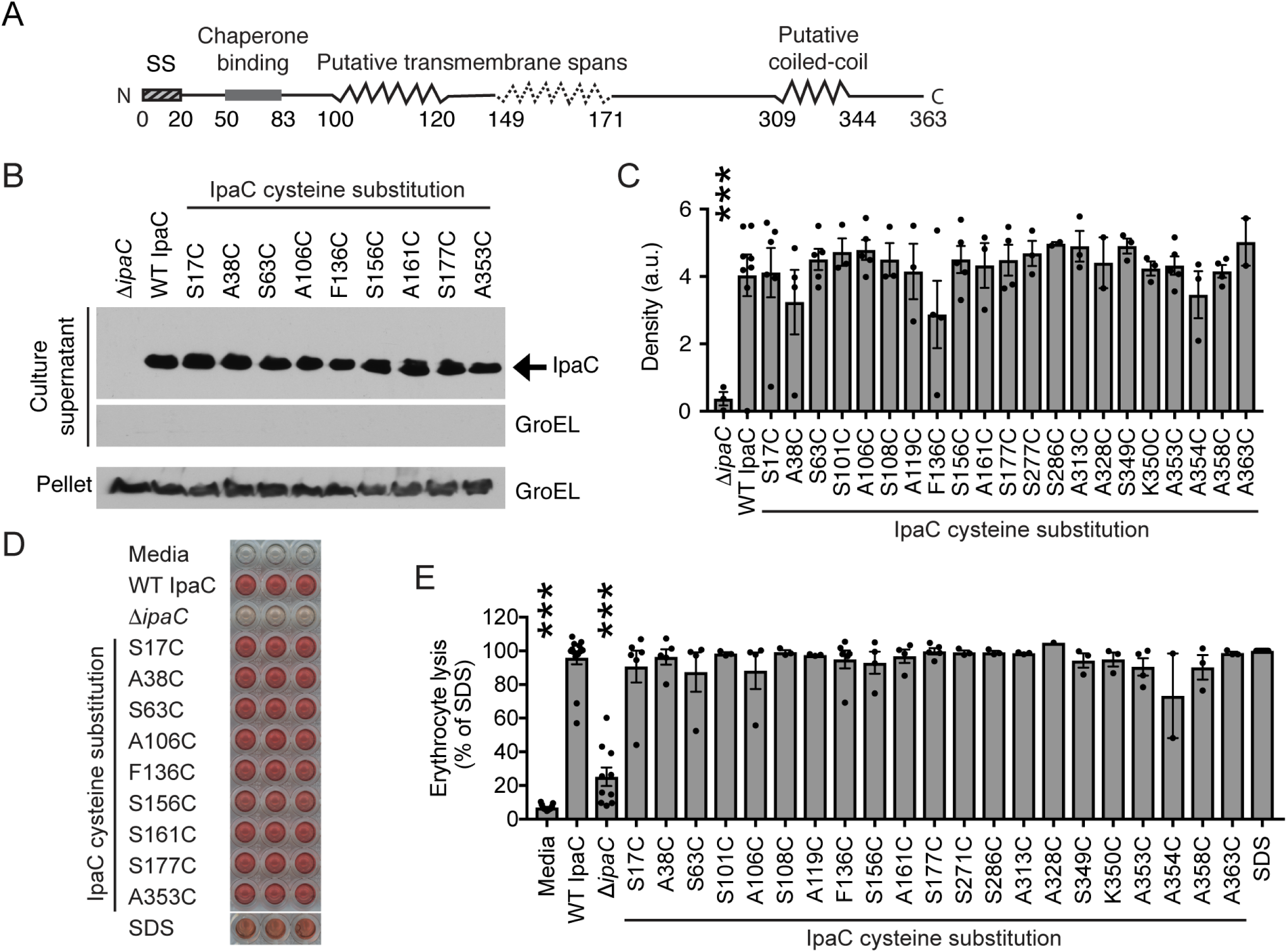
Cysteine substitution derivatives of IpaC support secretion and formation of translocon pores with efficiencies similar to WT IpaC. **(A)** Schematic showing the putative secondary structure of IpaC (7,9). Putative transmembrane span, residues 100-120 (solid zig-zag), predicted by *in silico* analyses and experimental data (7,9); second putative transmembrane span, residues 149-171 (dotted zig-zag), suggested by (10). **(B-C)** The efficiency by which each IpaC cysteine substitution derivative or WT IpaC, expressed in *S. flexneri* Δ*ipaC*, was secreted through the T3SS following activation of secretion by addition of Congo red to the medium. Soluble IpaC in the culture supernatant. **(B)** Representative western blots. IpaC and GroEL, bacterial cytosolic protein used as a control for bacterial lysis (culture supernatant) and for loading (pellet). **(C)** Densitometry analysis of bands for secreted IpaC from three independent experiments for each cysteine substitution derivative; mean ± SEM; dots represent individual experimental replicates. ***; p<0.001 (by ANOVA with Dunnett’s *post hoc* test comparing detectable IpaC secreted from *S. flexneri* Δ*ipaC* to *S. flexneri* Δ*ipaC* producing WT IpaC). Secretion of IpaC cysteine substitution derivatives was not significantly different from WT IpaC (by ANOVA with Dunnett’s *post hoc* test). **(D-E)** Erythrocytes were co-cultured with *S. flexneri* Δ*ipaC* strains producing IpaC containing a cysteine substitution derivative or WT IpaC; pore formation causes erythrocyte lysis and hemoglobin release. **(D)** Representative image showing hemoglobin released from erythrocytes. Each independent experiment was performed in triplicate. **(E)** The abundance of hemoglobin released was quantified by A_570_ from at least two independent experiments for each cysteine substitution mutant; mean ± SEM; dots represent independent experimental replicates. ***, p<0.001, compares hemoglobin release induced by indicated strains to hemoglobin release induced by each *S. flexneri* strain producing WT IpaC (by ANOVA with Dunnett’s *post hoc* test). Hemoglobin release induced by each strain producing cysteine substitution derivatives of IpaC was not statistically different from hemoglobin release induced by the strain producing WT IpaC (by ANOVA with Dunnett’s *post hoc* test).

### IpaC tolerates substitution with cysteine at numerous sites along the length of the protein

Selected IpaC residues were replaced with cysteine and the accessibility of these cysteines was determined by site-directed labeling with a chemical probe specific for the thiol group of the cysteine (12). Of note, native IpaC lacks cysteines. During infection, together with IpaB, IpaC is secreted through the T3SS, and embeds in and forms a pore in the plasma membrane. Type 3 substrates are secreted through the T3SS needle in an unfolded conformation, and proteins that are folded cannot traverse the needle (13). When the T3SS was artificially activated by incubation of bacteria with the dye Congo red (14), all IpaC cysteine substitution derivatives were readily secreted through the type 3 secretion needle into the culture supernatant (Fig. 1B-C), indicating that the cysteine substitutions did not disrupt or only minimally disrupted protein conformation in the bacterium.

The ability of the cysteine substitution derivatives of IpaC to support formation of translocon pores in plasma membranes was tested using the membranes of erythrocytes by co-culturing erythrocytes with strains of *S. flexneri* producing individual IpaC cysteine substitution derivatives. In this assay, the formation of translocon pores in the membrane of the erythrocytes leads to lysis of the erythrocyte and the release of hemoglobin. The abundance of hemoglobin released into the infection media, quantified spectrophotometrically, is an indicator of the efficiency of cell lysis. The absorbance of the supernatants was similar for strains of *S. flexneri* producing IpaC cysteine substitution derivatives and WT IpaC (Fig. 1D-E), which indicates that the IpaC derivatives support formation of translocon pores at an efficiency similar to that of WT IpaC.

To test the impact of the cysteine substitutions on the function of the translocon pores, translocon pore-related functions during infection of cells by *S. flexneri* were assessed. In a process termed docking (7,8,15), *Shigella* stably associates with eukaryotic cells via interactions between the T3SS needle and the translocon pore; IpaC is required for docking and subsequent effector translocation [Fig. 2A-D and (7)]. To test whether IpaC cysteine substitution derivatives supported docking, HeLa cells were infected with strains of *S. flexneri* producing an IpaC substitution derivative or WT IpaC, and the number of bacteria that stably associated with cells at 40 minutes of infection was quantified. *S. flexneri* strains producing an IpaC with a substituted cysteine docked to cells at similar numbers to *S. flexneri* producing WT IpaC (Fig. 2A and E).

**Figure 2:**
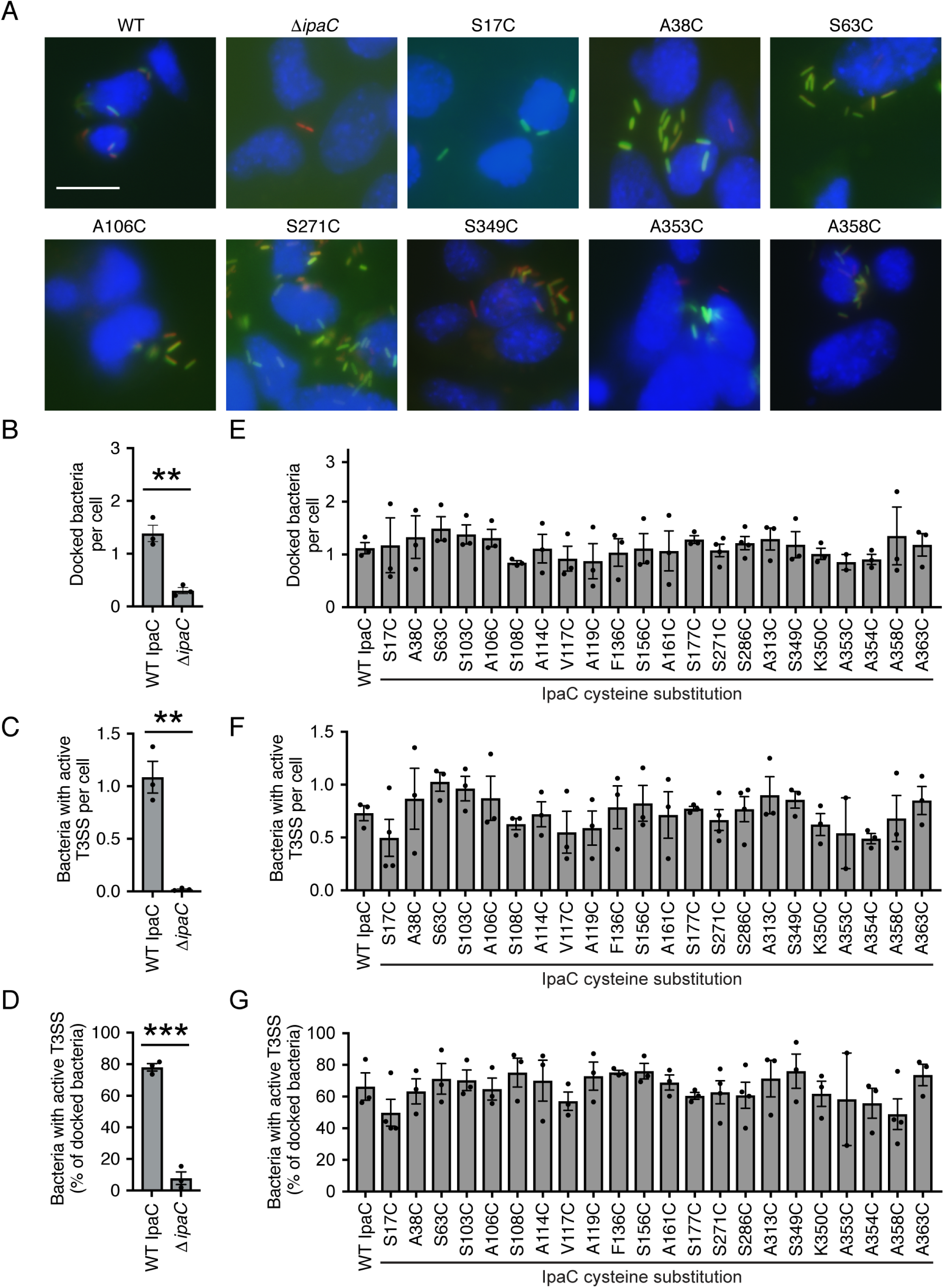
Cysteine substitution derivatives of IpaC support formation of translocon pores that function similar to those formed by WT IpaC. Stable docking and effector translocation upon infection of mouse embryonic fibroblasts cells with *S. flexneri* Δ*ipaC* strains producing IpaC derivatives containing a cysteine substitution derivative or WT IpaC. The secretion activity of the T3SS was measured using the TSAR reporter (7,16). **(A)** Representative images of infected cells. Blue, DAPI; red, mCherry (all bacteria); green, GFP (TSAR reporter; bacteria with active T3SS secretion). Scale bar, 20 μM. **(B and E)** Average number of bacteria that stably associated with cells (docked). RFP positive bacteria per cell. **(C and F)** Average number per cell of bacteria that produced GFP, indicating active T3SS secretion. GFP positive bacteria per cell. **(D and G)** The percentage of docked bacteria that produced GFP, indicating active T3SS secretion. GFP positive bacteria per RFP positive bacteria. Data are the mean ± SEM; dots represent independent experimental replicates. **, p<0.01; ***, p<0.001 (Student’s t-test). Stable docking of bacteria and secretion via the T3SS for each strain producing an IpaC cysteine substitution derivative was not statistically different from that of the strain producing WT IpaC (by ANOVA).

The ability of T3SS translocon pores formed by the IpaC cysteine substitution derivatives to activate effector secretion was monitored using a fluorescent transcription-based reporter of T3SS activity (TSAR, (7,16)); following the secretion of the translocon pore proteins and the bacterial effector protein OspD through the T3SS, the TSAR reporter produces GFP in the bacterium. The promoter for GFP is regulated by MxiE, which is activated upon secretion of OspD (16). *S. flexneri* producing cysteine substituted IpaC derivatives activated the T3SS to secrete effector proteins at efficiencies similar to WT IpaC (Figs. 2A, F, and G, and S1A). Both the absolute number per cell of bacteria with active secretion (Fig. 2A and F) and the percentage of docked bacteria with active secretion (Fig. 2A and G) were similar for *S. flexneri* producing IpaC containing cysteine substitutions and *S. flexneri* producing WT IpaC. These data demonstrate that cysteine substitution at the selected residues in IpaC did not adversely affect T3SS-mediated docking or secretion of effector proteins through the T3SS, including through the translocon pore.

*Shigella* invasion of cells requires the translocation of bacterial effector proteins through the translocon pore into the cytosol. The translocated effectors induce actin rearrangements, which in turn generate membrane ruffles that engulf the bacteria (17-19). The effect of cysteine substitutions in IpaC on *S. flexneri* invasion was tested by quantifying the numbers of bacteria internalized during infection of HeLa cells (7). The numbers of intracellular bacteria recovered were similar for HeLa cells infected with *S. flexneri* producing IpaC cysteine substitution derivatives and those infected with *S. flexneri* producing WT IpaC (Fig. S1B). In sum, substitution of selected IpaC residues with cysteine did not significantly alter either IpaC activity or T3SS function during *S. flexneri* infection of cells.

### Accessibility of IpaC residues in membrane-embedded *S. flexneri* translocon pores

To measure the accessibility of IpaC residues in translocon pores embedded in the plasma membrane of host cells, site-directed labeling by methoxypolyethelene glycol maleimide (PEG5000-maleimide) was performed with the library of IpaC cysteine substitution derivatives characterized above. Because PEG5000-maleimide is membrane impermeant (12) and is too big to pass entirely through the translocon pore (3,7), this approach specifically assesses the accessibility of the cysteine residue to the extracellular surface of the eukaryotic cell. Cysteine substitutions within the N-terminal domain (residues 1-99) of IpaC labeled efficiently with PEG5000-maleimide, as demonstrated by a distinct shift to a slower migrating position on SDS-PAGE (Fig. 3). Efficient labeling was also observed for one of the four residues tested within the putative transmembrane α-helix (residues 100-120) (Fig. 3); labeling at IpaC A106C was efficient, whereas labeling of the three other residues in the putative transmembrane α-helix that were tested (S101C, S108C, and A119C) was essentially undetectable. The labeling of IpaC cysteine substitutions C-terminal to the putative transmembrane α-helix was weak, except among substitutions of the 15 residues closest to the C-terminus (residues 349-363) (Fig. 3). All cysteine substitutions of IpaC, including the substitutions in the putative transmembrane α-helix and the substitutions C-terminal to it, were readily labeled by PEG5000-maleimide when the labeling was performed on IpaC in solution following secretion from bacteria *in vitro* (Fig. S2A-B), indicating that the lack of labeling observed during infection was a function of inaccessibility due to the membrane-embedded state rather than to fundamental inaccessibility of the residue in the protein *per se*. When labeling was performed following permeabilization of the plasma membrane, accessibility of IpaC residues near the C-terminus was similar to the accessibility of the residues in the N-terminal domain (Fig. S2C-D), which demonstrates that under non-permeabilizing conditions, the plasma membrane inhibited access of PEG5000-maleimide to the residues near the C-terminus. To test whether docking of the needle on the translocon pore inhibited labeling of residues near the C-terminus, we compared the accessibility of IpaC A358C, which is close to the C-terminus (R362), with that of S17C, a residue in the N-terminal domain, in the presence and absence of intermediate filaments. We previously showed that in the absence of intermediate filaments, docking does not occur but translocon pores form at wild-type levels (7). Since docking occurs through interactions of the needle tip complex with the pore (15), in the absence of intermediate filaments, pores form without a needle tip complex associated with the pore. The accessibility of A358C was similar in the presence and absence of intermediate filaments (Fig. S3), and the accessibility of S17C was greater than the accessibility of A358C and was also independent of intermediate filaments (Fig. S3). These data show that the engagement of the T3SS needle and/or tip complex with the translocon pore does not alter the accessibility of residues near the C-terminus of IpaC. Altogether, these results indicate that, whereas IpaC sequences immediately C-terminal to the putative transmembrane α-helix are intracellular, some or all of the 15 residues closest to the C-terminus of IpaC are partially accessible from the extracellular surface of the plasma membrane.

**Figure 3:**
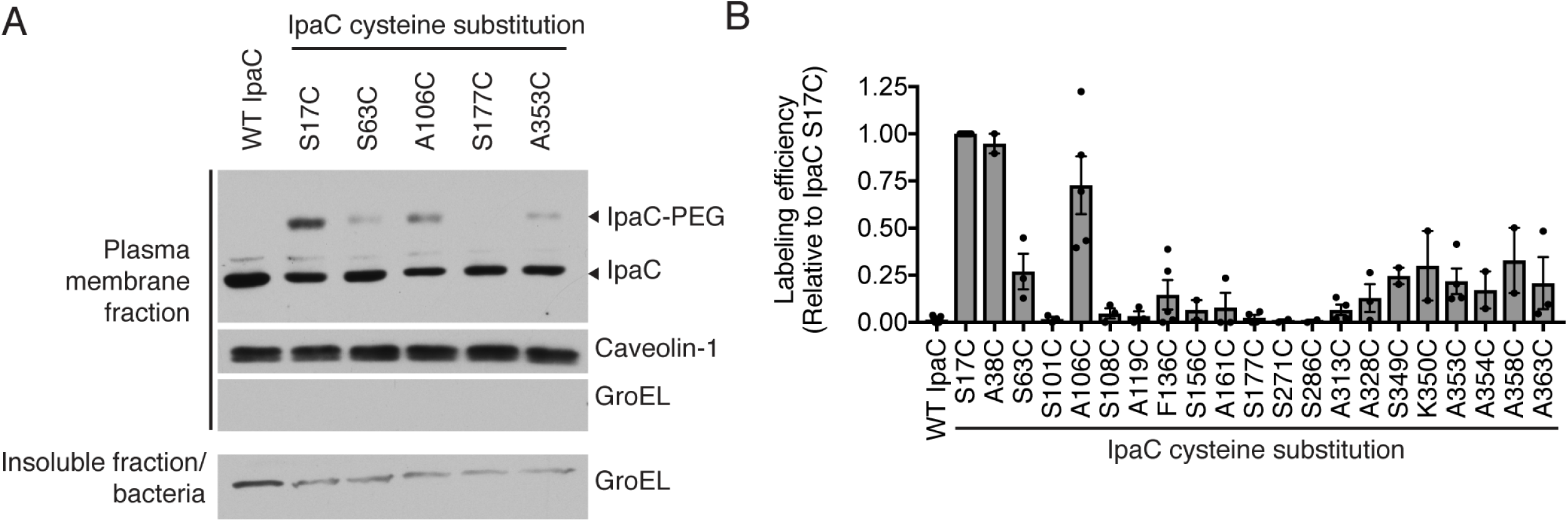
Topology of membrane-embedded IpaC upon native delivery of the translocon pore. Accessibility of membrane-embedded IpaC to labeling with PEG5000-maleimide upon infection of HeLa cells with *S. flexneri*. *S. flexneri* Δ*ipaC* strains producing IpaC cysteine substitution mutants or WT IpaC. **(A)** Gel shift of PEG5000-maleimide labeled IpaC in the plasma membrane fraction of infected HeLa cells. Representative western blots. IpaC-PEG, IpaC labeled with PEG5000-maleimide; IpaC, unlabeled IpaC; caveolin-1, eukaryotic plasma membrane protein; GroEL, bacterial cytosolic protein. **(B)** Relative accessibility of IpaC cysteine substitutions. Densitometry analysis of IpaC-PEG5000 bands from three independent experiments for each cysteine substitution derivative; mean ± SEM; dots represent independent experimental replicates.

### Accessibility of *Salmonella* translocon pore protein SipC in membrane-embedded *Salmonella* translocon pores

The *Shigella* T3SS is closely related to the *Salmonella* T3SS (1). The translocon pore of *Salmonella* is composed of SipB and SipC, with SipB being homologous to IpaB and SipC being homologous to IpaC. Alignment of SipC and IpaC by Clustal Omega (20-22) showed that 33% of IpaC residues are identical to those in SipC and another 26% of IpaC residues retain similar functional properties to those in SipC, together yielding 52% amino acid similarity of IpaC to SipC (Fig. S4). Both proteins are required for translocon pore formation, docking, and effector protein secretion; however, transcomplementation experiments show that the two proteins do not function equivalently in all aspects of infection. Whereas *S.* Typhimurium are predominantly found within vacuoles, complementation of a *S.* Typhimurium *sipC* mutant with *Shigella ipaC* enables bacterial escape from the vacuole (23), which indicates that SipC and IpaC are sufficient to direct the bacterium to distinct intracellular niches (23).

To explore whether the observed functional differences between IpaC and SipC were associated with differences in protein topology in the plasma membrane, we compared the accessibility of SipC residues with that of IpaC residues. Like IpaC, native SipC lacks cysteines. The putative secondary structure of SipC is very similar to that of IpaC, with a putative N-terminal extracellular domain (residues 1-119), a putative transmembrane α-helix (residues 120-140), and C-terminal to the transmembrane α-helix, a putative coiled-coil domain (residues 293-320) (Fig 4A). The second putative transmembrane span identified in IpaC was not clearly delineated by *in silico* analyses of SipC. We substituted cysteines at residues of SipC that were analogous to those chosen for substitution in IpaC (Fig. 4B-C). During *S.* Typhimurium infection of HeLa cells, the accessibility of these residues from the extracellular surface of the plasma membrane was tested using the same method used for IpaC, site-directed labeling with PEG5000-maleimide. As observed for IpaC, substitutions in the N-terminal domain of SipC labeled efficiently (Fig. 4D-E). In contrast to our findings for IpaC, labeling was not observed for any other SipC substitution tested (Fig. 4D-E); this included SipC A126C, a cysteine substitution at a residue within the SipC putative transmembrane α-helix that is analogous to IpaC A106C, which labeled efficiently with PEG5000-maleimide, and cysteine substitutions at residues within the C-terminal 15 amino acids of SipC, which in IpaC labeled with intermediate efficiency.

**Figure 4:**
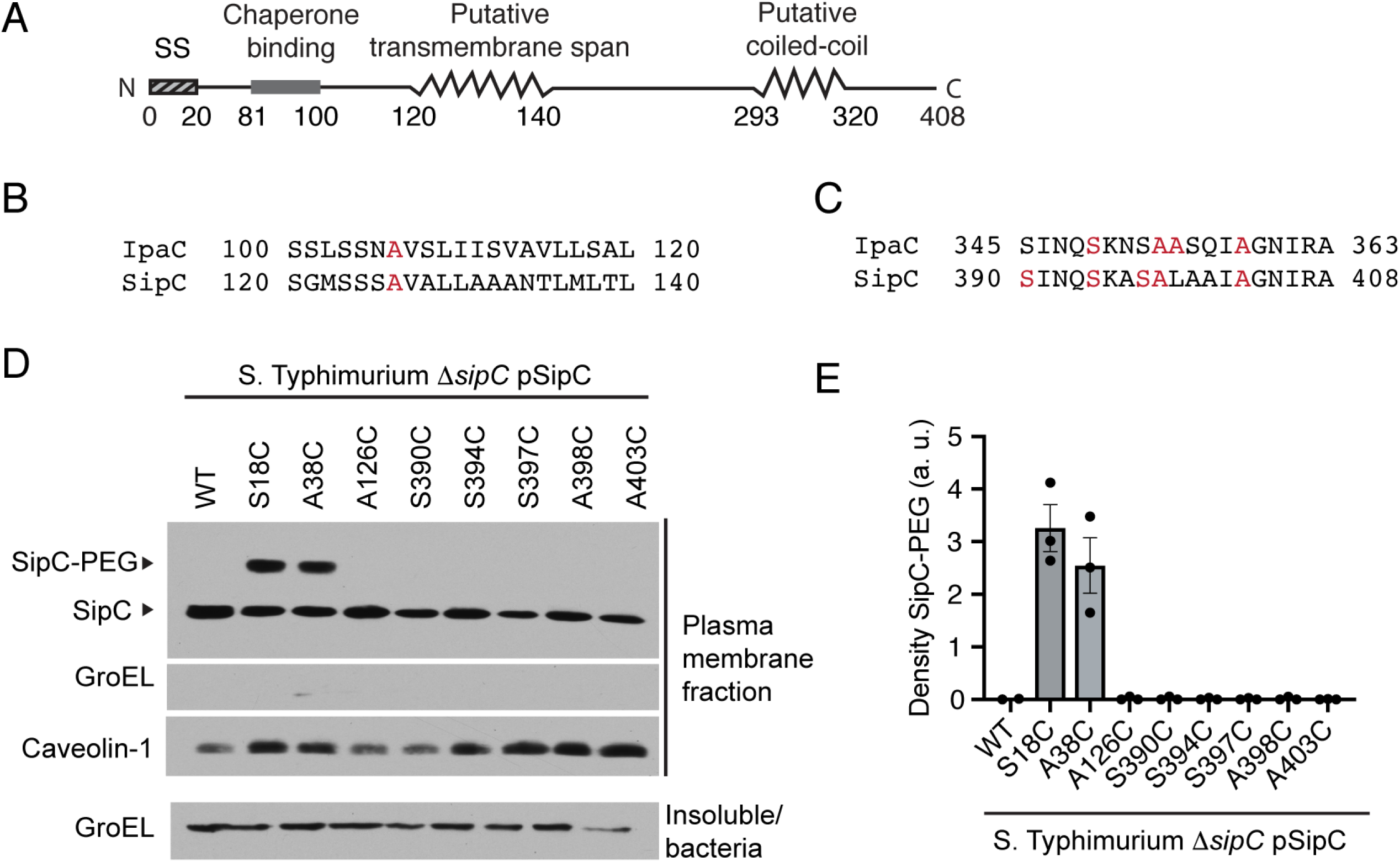
SipC is inserted in the plasma membrane with the N-terminal domain extracellular and the C-terminal domain intracellular. **(A)** Schematic showing the putative secondary structure of SipC. **(B-C)** Alignment by Clustal Omega of IpaC and SipC transmembrane α-helix (B) and C-terminal 19 amino acids (C). **(D-E)** Accessibility of membrane-embedded SipC residues to the extracellular surface of the plasma membrane. Labeling of SipC with PEG5000-maleimide upon infection of HeLa cells with *S.* Typhimurium *ΔsipC* producing SipC cysteine substitution derivatives or WT SipC. **(D)** Gel shift of PEG5000-maleimide labeled SipC in the plasma membrane fraction of infected HeLa cells. Representative western blots. SipC-PEG, SipC labeled with PEG5000-maleimide; SipC, unlabeled SipC; caveolin-1, eukaryotic plasma membrane protein; GroEL, bacterial cytosolic protein. **(E)** Relative accessibility of SipC cysteine substitutions. Densitometry analysis of SipC-PEG5000 bands from three independent experiments for each cysteine substitution derivative. Mean ± SEM; dots represent independent experimental replicates.

As for IpaC, SipC tolerated substitution with cysteine at numerous sites along the length of the protein. The SipC cysteine derivatives were produced in *Salmonella* at levels similar to WT SipC (Fig. S5A). Moreover, each cysteine substitution derivative supported *S.* Typhimurium docking to HeLa cells (Fig. S5B). Thus, the differences in labeling observed for cysteine substitutions of IpaC compared to cysteine substitutions of SipC were not due to altered function of the SipC derivatives.

These data suggest that the overall topologies of plasma membrane-embedded SipC and plasma membrane-embedded IpaC are similar. For each protein, the N-terminal domain is extracellular, a single transmembrane α-helix is present, and the C-terminus is intracellular (Fig. 5). Despite the similarities in overall topology, the differences observed in the accessibility of substitutions close to the C-termini of the two proteins suggest that subtle topological or structural differences exist in the transmembrane α-helix and C-terminal regions. The differences we observe between the two proteins in the context of plasma membrane-embedded translocon pores may be relevant to functional differences observed for *Shigella* and *Salmonella* translocon pores.

**Figure 5:**
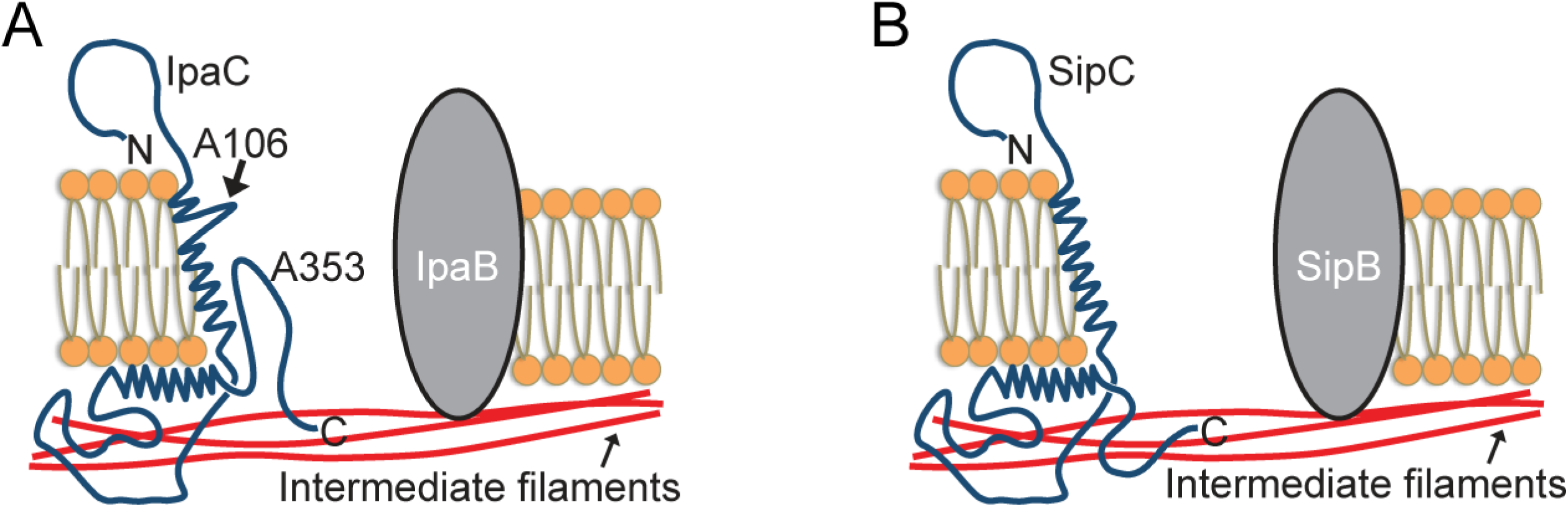
Topological models of plasma membrane-embedded *Shigella* IpaC and plasma membrane-embedded *Salmonella* SipC. **(A)** Topological model of the *Shigella* translocon pore protein IpaC. The N-terminal domain of IpaC is situated on the extracellular side of the membrane. A single transmembrane α-helix is present. IpaC A106 is positioned such that it is accessible to the lumen of the pore. The region C-terminal to the transmembrane α-helix, including the coiled-coil domain, is situated on the cytosolic side of the membrane. IpaC A353, which is among the C-terminal 15 residues, is present within a loop of IpaC that is positioned within the pore lumen and accessible to the extracellular side of the plasma membrane. **(B)** Topological model of the *Salmonella* translocon pore protein SipC. The N-terminal domain of SipC is situated on extracellular side of the plasma membrane. A single transmembrane α-helix is present. The C-terminal domain is situated on the cytosolic side of the plasma membrane.

## Discussion

The T3SS is essential for the pathogenesis of many important human bacterial pathogens, and the translocon pore is essential for T3SS function. To generate insights into how the translocon pore participates in the signaling that activates secretion of effector proteins, we defined the topology of IpaC in the membrane of human-derived cells. To our knowledge, this represents the first detailed topological analysis of a type 3 translocon pore protein following its native delivery to the plasma membrane. We show that the N-terminal region of IpaC is extracellular and the C-terminal region of IpaC is in the host cytosol. Moreover, we observed a similar overall topology for SipC, the IpaC homolog in *Salmonella*. The observations that the N-terminal region of IpaC is located on the extracellular side of the plasma membrane is consistent with previous observations investigating the interactions of recombinant IpaC with the membrane (9,10).

The labeling of A106C in the transmembrane span indicates that the outer portion of the pore channel is at least 4.4 nm wide, the approximate size of PEG5000-maleimide. Moreover, the lack of labeling for other residues in the transmembrane α-helix of IpaC suggest that towards the cytosolic side of the plasma membrane, either the lumen of the pore becomes narrower, such that the PEG5000-maleimide is precluded from accessing the IpaC residues, and/or that other IpaC residues do not line the pore interior. If the latter were true, it could indicate that IpaB residues line the bulk of the channel.

C-terminal to the predicted transmembrane α-helix are sequences without predicted secondary structure (residues 121-308) and a predicted coiled-coil domain (residues 309-344). The minimal accessibility to PEG5000-maleimide of IpaC cysteine substitutions within these sequences strongly suggests that these sequences, including the coiled-coil domain, lie within the cytosol and that residues 100-120 constitute a transmembrane α-helix and the only transmembrane α-helix in the protein. By extension, these data suggest that all of IpaC C-terminal to this transmembrane α-helix lies on the cytosolic side of the plasma membrane (Fig. 5A). Additional attempts to directly assess the location of the C-terminal region of IpaC, including limited proteolysis, incorporation of enzymatic cleavage sites that would specifically function within the intracellular environment, and immunolabeling with monoclonal antibodies specific for C-terminal or N-terminal sequences, were uninformative.

Cysteine substitutions in the C-terminal 15 amino acids of IpaC were weakly but reproducibly accessible from the outside of the cell, suggesting that they are present within the pore lumen. Consistent with this possibility, crosslinking studies showed that residues in the C-terminal region of the *Pseudomonas* translocon pore protein PopD, a homolog of IpaC, interact with the *Pseudomonas* T3SS tip complex protein (15), which docks onto the extracellular face of the translocon pore. Of note, atomic force microcopy studies suggest that the lumen of the *E. coli* translocon pore is shaped like a funnel, with the narrower portion of the funnel closer to the cytosolic side of the plasma membrane (24). If the *Shigella* translocon pore is shaped similarly to the *E. coli* translocon pore, the detection of PEG5000-maleimide labeling of cysteine substitutions near the C-terminus of IpaC could be consistent with these residues looping back from the cytosolic side of the plasma membrane into the pore lumen where they might be positioned to engage the tip complex. If this is the case, these C-terminal domain residues might contribute directly to the narrowest portion of the funnel shape of the pore.

Our data support a model of the native topology of IpaC in mammalian plasma membranes in which the N-terminal domain of IpaC is extracellular, a single transmembrane α-helix crosses the plasma membrane, the region C-terminal to the transmembrane α-helix is present within the cytosol, and the C-terminal 15 amino acids of IpaC re-enter the pore interior (Fig 5A).

The overall topology of SipC and IpaC was similar, with the N-terminus of SipC extracellular and the C-terminus intracellular (Fig. 5B). Both IpaC and SipC support bacterial docking to cells (7,8). Our models place the N-terminal regions of these proteins at the cell surface where these sequences would be positioned adjacent to the T3SS needle as it docks on the extracellular face of the translocon pore. This proximity would facilitate the involvement of the N-terminal domains of these proteins in docking and activation of effector secretion. Moreover, for both *Shigella* IpaC and *Salmonella* SipC, an interaction with intermediate filaments is required for bacterial docking (7). For IpaC, we previously demonstrated that sequences adjacent to the protein’s C-terminus are required for this interaction with intermediate filaments (7). Our models for IpaC and SipC place the extreme C-terminal regions of these proteins in the cytosol of the eukaryotic cell, adjacent to the intermediate filaments of the cell cortex with which they interact.

The similarity between the overall topology of IpaC and that of SipC suggests that the mechanism(s) required to deliver the two proteins into the membrane, multimerize into the pore, interact with the lipid membrane, support effector translocation, and generate signals to activate secretion may be similar. However, in contrast to the accessibility of some cysteine substitutions in the C-terminal region of IpaC, substitutions near the C-terminus of SipC were completely inaccessible. This indicated that multiple topologies of SipC do not occur at an efficiency that was detectable in this assay and, by extension, that the weak labeling observed at IpaC substitutions near the C-terminus are unlikely to be the result of a minority population of membrane-embedded IpaC having an alternate topology. In addition, whereas the two proteins are homologs, they display functional differences. When IpaC and SipC are swapped, IpaC partially complements the loss of SipC but SipC is unable to complement the loss of IpaC (23,25), and IpaC production in *S.* Typhimurium heterologously enables the bacterium lyse the vacuolar membrane (23). Together, our findings raise the possibility that the observed subtle differences in the topological arrangements of SipC and IpaC may contribute to their functional differences during bacterial infection. We speculate that the greater accessibility in IpaC of residues within the transmembrane α-helix and near the C-terminus reflects a more open channel in *Shigella* translocon pores than in *Salmonella* translocon pores. Such a confromation may be associated with lysis of the vacuolar membrane in the case of *Shigella* and maintenance of the vacuolar membrane in the case of *Salmonella*. Further investigations into translocon pore structure will inform the molecular mechanisms that drive the functional differences among the translocon pore proteins.

## Materials and Methods

### Bacterial culture

Strains used in this study are described in Table 1. The *S. flexneri* wild type strain was serotype 2a strain 2457T (26), and all mutants were isogenic derivatives of it. *S. flexneri* strains were cultured in trypticase soy broth (Benton Dickenson) at 37°C. *ipaC* derivatives were encoded on the plasmid pBAD33 and their expression was driven from the pBAD promoter, induced with 1.2% arabinose. The sequences of primers used for PCR and sequencing are available from the authors upon request. The wild type *Salmonella enterica* serovar Typhimurium strain was SL1344, and all *S.* Typhimurium mutants used in this study were isogenic derivatives of it. *Salmonella* strains were cultured in trypticase soy broth at 37°C. *sipC* derivatives were encoded on the plasmid pBAD18 and their expression was driven from the pBAD promoter, induced with 1.2% arabinose.

**Table 1:**
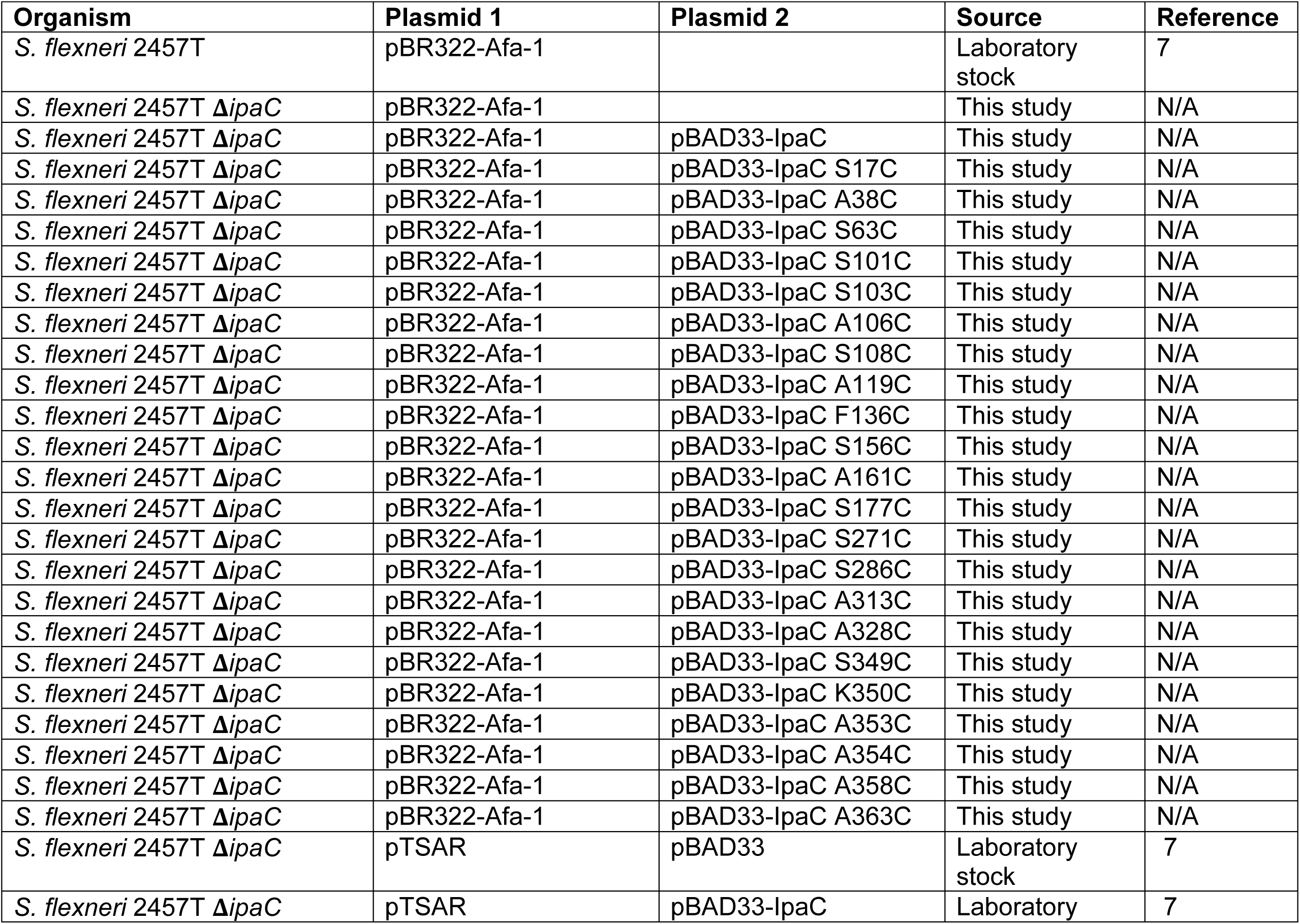
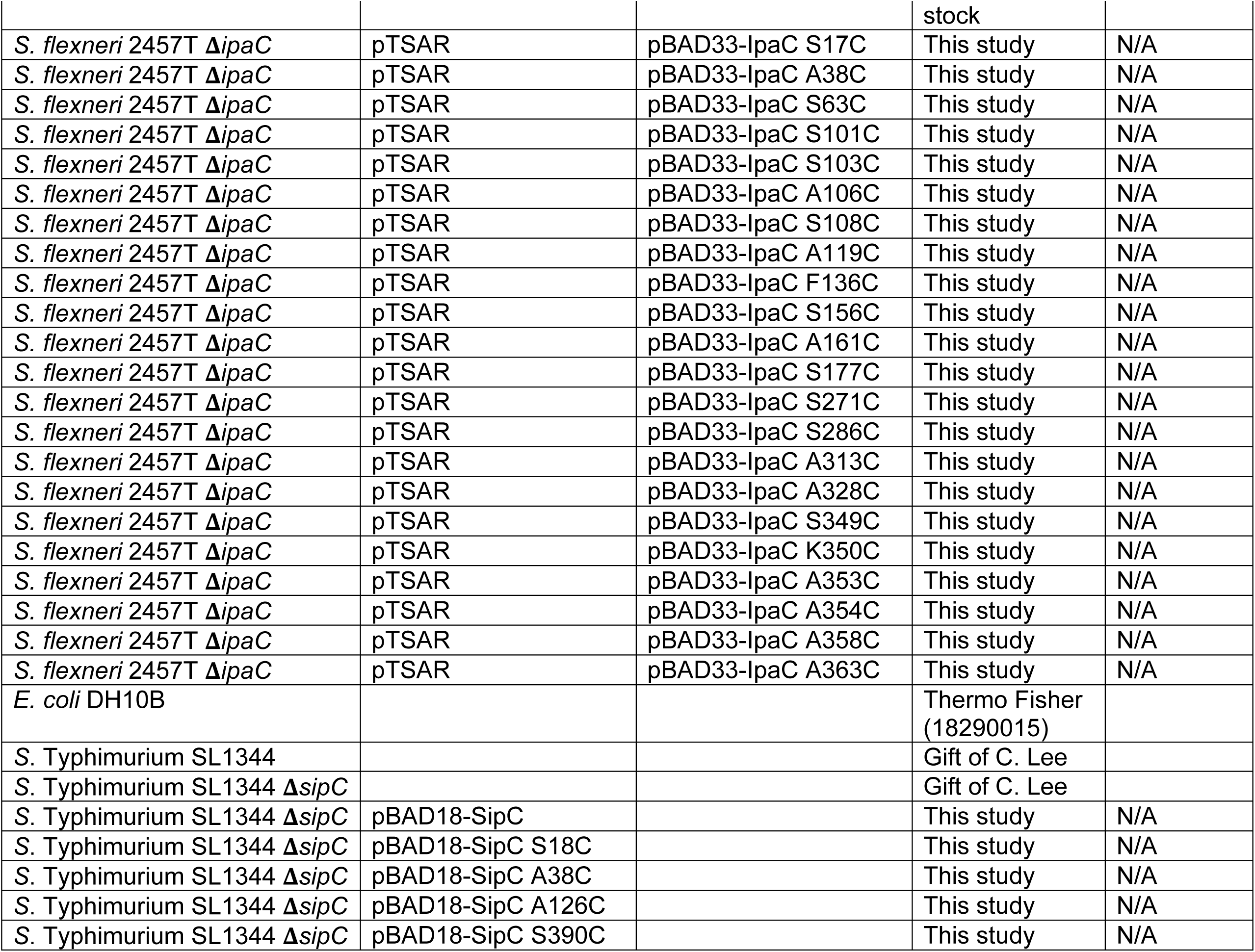
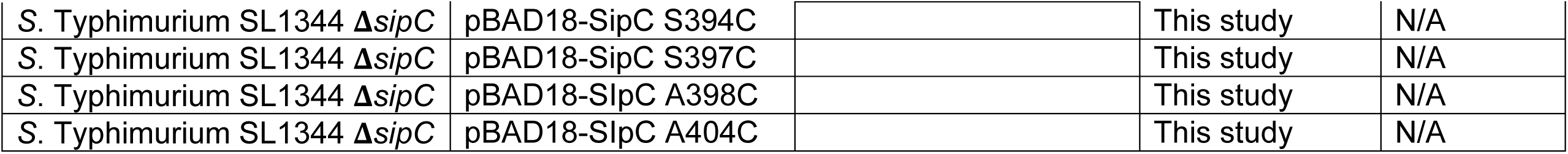
Strains used in this study.

### Cell culture

Mouse embryonic fibroblasts were kindly provided by Victor Faundez (Emory), and HeLa (CCL2) cells were obtained from ATCC. All cells were cultured in Dulbecco’s Modified Eagles Media (DMEM) supplemented with 0.45% glucose and 10% heat-inactivated fetal bovine serum (FBS); they were maintained at 37°C in humidified air containing 5% CO_2_. All cells in the laboratory are periodically tested for mycoplasma; cells that test positive are treated or discarded.

### Secretion assays

To assess the ability of the type 3 secretion system to secrete proteins in the absence of eukaryotic cells, the T3SS was artificially activated by addition of Congo red (7,14,27). Overnight cultures were back diluted, induced where appropriate with 1.2% arabinose, and cultured for 2 hours at 37°C. Bacterial cultures were normalized by OD_600_, resuspended in PBS containing 1.2% arabinose, where appropriate, and 10 μM Congo red, and incubated in a water bath at 37°C for 45 min. The bacteria were removed by centrifugation, and the resulting supernatant containing the secreted proteins was collected.

### Erythrocyte lysis assay

Pore formation in sheep erythrocyte membranes was monitored by assessing the efficiency of erythrocyte lysis (3,7). Briefly, 10^8^ erythrocytes were washed with saline and infected with *S. flexneri* at a multiplicity of infection (MOI) of 25 in 30 mM Tris, pH 7.5. The bacteria were centrifuged onto the erythrocytes at 2000 *g* for 10 minutes at 25°C. Bacteria and erythrocytes were cultured together for 30 minutes at 37°C in humidified air with 5% CO_2_. The bacteria and erythrocyte co-cultures were mixed by pipetting and then centrifuged again at 2,000 *g* for 10 minutes at 25°C. As a positive control for lysis, an aliquot of uninfected erythrocytes was treated with 0.02% SDS and then centrifuged. The supernatants were collected, and the abundance of hemoglobin released was measured spectrophotometrically at A_570_ using a Wallac 1,420 Victor^2^ (Perkin Elmer).

### Translocation and docking

Mouse embryonic fibroblasts were seeded at 3×10^5^ cells per well on coverslips in a 6-well plate. The cells were infected with exponential phase *S. flexneri* (18,28) harboring the TSAR reporter (16) at a MOI of 200. The TSAR reporter, which expresses GFP when the bacterial effector OspD is secreted through the T3SS (16), is an indicator of active T3SS secretion. Bacteria were centrifuged onto cells at 800 *g* for 10 minutes at 25°C. The bacteria and cells were co-cultured for an additional 30 minutes at 37°C in humidified air with 5% CO_2_. Cells were washed three times with PBS, fixed with 3.7% paraformaldehyde for 20 minutes at 25°C, and washed again with PBS. DNA was stained with Hoechst. Four random microscopic fields per experimental condition were imaged by fluorescent microscopy using a Nikon Eclipse TE-300 with appropriate filters. All bacteria produced mCherry, and bacteria in which T3SS secretion was activated produced GFP (16). For each condition of each independent experiment, 20-250 eukaryotic cells and 20-210 bacteria were analyzed.

### Quantification of intracellular bacteria

Hela cells were seeded at 1.5×10^4^ cells per well in 96-well plates the day prior to infection. *S. flexneri* strains were grown as described above and added to cell monolayers in Hanks Balanced Salt Solution (HBSS) supplemented with 1.2% arabinose at a MOI of 100. Bacteria were centrifuged onto cells for 10 min at 2,000 rpm and incubated at 37°C with 5% CO_2_ for 20 minutes. Bacteria that did not invade were removed by washing with HBSS. The HeLa cells and bacteria were co-cultured for an additional hour at 37°C with fresh HBSS containing 25 µg/ml gentamicin, which kills extracellular but not intracellular bacteria. Infected monolayers were then washed with phosphate buffered saline and lysed in PBS containing 0.5% Triton X-100 to release the intracellular bacteria. Lysates containing the bacteria and plated on agar plates with appropriate antibiotics in order to quantify the number of intracellular bacteria.

### Labeling of cysteines with PEG5000-maleimide

Individual IpaC or SipC amino acids were replaced with cysteines; native IpaC and native SipC both lack cysteines. For the analysis of the efficiency of PEG5000-maleimide labeling of soluble IpaC, 2.5 mM PEG5000-maleimide was added to supernatants collected following induction of T3SS secretion with Congo red. The supernatant was incubated with PEG5000-maleimide at 30°C for 30 min. Samples were analyzed by SDS-PAGE and western blot; the efficiency of labeling was monitored by assessing the gel shift of IpaC.

For the analysis of cysteine labeling during infection, HeLa cells were seeded at 4×10^5^ cells per well in a 6-well plate. For each strain tested, 8×10^5^ HeLa cells or 2.4×10^6^ MEFs were used per experimental condition. Cells were washed once with 50 mM Tris, pH 7.4, supplemented with 150 mM NaCl and 1.2% arabinose. To enhance the efficiency of translocon pore insertion into the HeLa cell membrane, bacteria expressed the *E. coli* adhesion Afa-1 (7,29). HeLa cells were infected with *S. flexneri* Δ*ipaC* producing individual IpaC cysteine substitution derivatives and Afa-1 or with *S.* Typhimurium Δ*sipC* producing individual SipC cysteine substitution derivatives at a multiplicity of infection (MOI) of 200 in 50 mM Tris, pH 7.4, supplemented with 150 mM NaCl, 1.2% arabinose, and 2.5 mM PEG5000-maleimide. The bacteria were centrifuged onto the cells at 800 *g* for 10 minutes at 25°C and incubated at 37°C in humidified air with 5% CO_2_ for 20 min. Cells were then harvested and plasma membrane-enriched fractions were isolated, as done previously (7,30). Briefly, the cells were washed three times with ice-cold 50 mM Tris, pH 7.4, and scrapped in 50 mM Tris, pH 7.4, containing protease inhibitors (Protease inhibitor cocktail, complete mini-EDTA free, Roche). Scrapped cells were pelleted at 3,000 *g* for 3 minutes at 25°C. The pelleted cells were resuspended in 50 mM Tris, pH 7.4, containing protease inhibitors and 0.2% saponin, and were incubated on ice for 20 min. For experiments testing the accessibility of IpaC residues in the presence of membrane permeabilization, PEG5000-maleimide was added in the presence of 0.2% saponin and not included during the infection. The suspension was centrifuged at 21,000 *g* for 30 minutes at 4°C. The supernatant, which contains the cytosol fraction, was decanted into a fresh tube. The pellet was resuspended in 50 mM Tris, pH 7.4, containing protease inhibitors and 0.5% Triton X-100, incubated on ice for 30 minutes, and centrifuged at 21,000 *g* for 15 minutes at 4°C. The supernatant from this spin contained the membrane fraction, and the pellet consisted of the detergent insoluble fraction, which included intact bacteria. The efficiency of PEG5000-maleimide labeling was monitored by assessing the gel shift of IpaC or SipC by western blot. Antibodies used for western blots: rabbit anti-IpaC (gift from Wendy Picking, 1:10,000), rabbit anti-GroEL (Sigma, G6352, 1:1,000,000), rabbit anti-Caveolin-1 (Sigma, cat # C4490, 1:1000), chicken anti-vimentin (BioLegend, cat # 919101, 1:5,000), mouse anti-SipC (Gift from Jorge Galán, 1:10,000), goat anti-rabbit conjugated with HRP (Jackson ImmunoResearch, cat # 115-035-003, 1:5,000), goat anti-mouse conjugated with HRP (Jackson ImmunoResearch, cat # 111-035-003, 1:5,000), donkey anti-chicken conjugated with HRP (Jackson ImmunoResearch, cat # 703-035-155, 1:5,000).

### Statistical analysis

Except where specifically noted, all data are from three independent experiments, and the means ± the standard error of the means (SEM) are presented. Dots presented within graphs represent independent experimental replicates. The means between groups were compared by a one-way analysis of variance (ANOVA) using GraphPad Prism 8 (GraphPad Software, Inc.). Alignment of IpaC and SipC was performed using Clustal Omega (20-22). Signal from western blots was captured by film, film was digitized using an Epson Perfection 4990 PHOTO scanner, and the density of bands was determined using ImageJ (National Institutes of Health).

## Acknowledgments

We thank Jorge Galán, Wendy Picking, and Cathy Lee for reagents and helpful discussions, Austin Hachey for technical assistance, and Daniel Kahne, D. Borden Lacy, Benjamin Spiller, Cammie Lesser, Amy Barczak, and the members of their laboratories for helpful discussions. This work was funded by NIH grant AI081724 to MBG, and by NIH grants AI007061 and AI114162, the Massachusetts General Hospital Executive Committee on Research Tosteson Award, and the Charles A. King Trust Postdoctoral Research Fellowship Program, Bank of America, N.A., Co-Trustees, to BCR.

## Conflicts of interest

The authors declare that they have no conflicts of interest with the contents of this article.

*The content is solely the responsibility of the authors and does not necessarily represent the official views of the National Institutes of Health.

## Supplemental files

**Supplement Number 1: Supplemental Figure Legends.**

**Supplement Number 2: Supplemental Figure 1. Cysteine substitution derivatives of IpaC support the activation of T3SS secretion and bacterial invasion. (A)** Secretion activity as GFP signal from TSAR reporter. Corresponding merged images are those shown in Fig. 2A. Blue, DAPI; red, mCherry (all bacteria); green, GFP (TSAR reporter; bacteria with active T3SS secretion). **(B)** Efficiency of *S. flexneri* invasion of HeLa cells, as measured by gentamicin protection assay. *S. flexneri* Δ*ipaC* strains producing IpaC derivatives containing a cysteine substitution derivative or WT IpaC. Data are the mean ± SEM of at least two independent experiments per strain, dots represent independent experimental replicates. *, p<0.05 (ANOVA with Dunnetts’ *post hoc* test) comparing the efficiency of invasion of *S. flexneri* Δ*ipaC* to *S. flexneri* Δ*ipaC* producing WT IpaC; the efficiency of invasion by each strain producing an IpaC cysteine substitution derivative was not significantly different from that of the strain producing WT IpaC.

**Supplement Number 3: Supplemental Figure 2. Permeabilization of the plasma membrane increased accessibility of IpaC residues near the C-terminus. (A-B)** Cysteine substitutions in IpaC are accessible *in vitro*. The accessibility of cysteine substitutions in soluble IpaC as assessed by reactivity with PEG5000-maleimide. Soluble IpaC from culture supernatants following activation of secretion by addition of Congo red to the medium. **(A)** Western blot for IpaC and GroEL. IpaC-PEG, IpaC conjugated with PEG5000-maleimide; IpaC, unlabeled IpaC; GroEL, bacterial cytosolic protein used as a control for bacterial lysis. **(B)** Densitometry analysis of IpaC-PEG5000 bands from independent replicates for each cysteine substitution mutant; mean ± SEM; dots represent individual experimental replicates. **(C-D)** During infection, following plasma membrane permeabilization, accessibility of IpaC residues near the C-terminus is similar to that of IpaC residues in the N-terminal domain. PEG-maleimide labeling of selected cysteine substitution derivatives after plasma membrane permeabilization of infected HeLa cells. **(C)** Gel shift of PEG5000-maleimide labeled IpaC in the plasma membrane fraction of infected HeLa cells. Representative western blot. IpaC-PEG, IpaC labeled with PEG5000-maleimide; IpaC, unlabeled IpaC; caveolin-1, eukaryotic plasma membrane protein; GroEL, bacterial cytosolic protein. **(D)** Densitometry analysis of IpaC-PEG5000 bands from four independent experiments for each cysteine substitution derivative; mean ± SEM; dots represent independent experimental replicates.

**Supplement Number 4: Supplemental Figure 3. Accessibility of PEG5000-maleimide to IpaC residues near the C-terminus was not altered by docking of the T3SS needle tip complex.** The accessibility of membrane-embedded IpaC S17C and IpaC A358C were tested in mouse embryonic fibroblasts (MEFs) derived from vimentin knockout mice (Vim^−/−^) or wild-type mice (Vim^+/+^). **(A)** Gel shift of PEG5000-maleimide labeled IpaC in the plasma membrane fraction of infected MEFs. Representative western blots. IpaC-PEG, IpaC labeled with PEG5000-maleimide; IpaC, unlabeled IpaC; caveolin-1, eukaryotic plasma membrane protein; GroEL, bacterial cytosolic protein. **(B)** Relative accessibility of IpaC cysteine substitutions. Densitometry analysis of IpaC-PEG5000 bands from three independent experiments for each cysteine substitution derivative; mean ± SEM; dots represent independent experimental replicates. N.S., not significant (by two-way ANOVA followed by a Sidak *post hoc* test). Although reduced in each experiment, labeling of S17C in Vim^−/−^ cells was not statistically different from labeling of S17C in Vim^+/+^ cells.

**Supplement Number 5: Supplemental Figure 4. Alignment of IpaC and SipC amino acid sequences.** Alignment performed by Clustal Omega. *, identical residues;:, residues with similar functional properties.

**Supplement Number 6: Supplemental Figure 5. SipC cysteine substitutions are produced in *S*. Typhimurium and support *S*. Typhimurium docking to cells. (A)** Protein production in *S*. Typhimurium Δ*sipC* is similar for plasmid-borne SipC cysteine substitution derivatives and plasmid-borne WT SipC. Western blot is representative of two independent experiments. **(B)** Docking on HeLa cells of *S*. Typhimurium Δ*sipC* strains producing SipC cysteine substitution derivatives or WT SipC. Images are representative of two independent experiments. Blue, DNA; green, *S*. Typhimurium.

